# CAR-T Cells with Phytohemagglutinin (PHA) Provide Anti-Cancer Capacity with Better Proliferation, Rejuvenated Effector Memory, and Reduced Exhausted T Cell Frequencies

**DOI:** 10.1101/2022.10.25.509664

**Authors:** Gamze Gulden, Berranur Sert, Tarik Teymur, Yasin Ay, Nulifer Neslihan Tiryaki, Ercüment Ovalı, Nevzat Tarhan, Cihan Tastan

**Author notes:** Equally contributed first authors (Gamze Gulden, Berranur Sert). **Author Contributions:** G.G., B.S, T.T., Y.A, N.N.T., and C.T. designed, completed, and/or analyzed preclinical experiments. G.G., B.S, T.T., Y.A, N.N.T., and C.T. wrote the manuscript and E.O. and N.T. contributed to the editing of the manuscript. C.T. led the project and contributed to the design and interpretation of the data.

## Abstract

The development of genetic modification techniques has led to the opening of a new era in cancer treatments that have been limited to conventional treatments such as chemotherapy. Since not only cancerous cells but also healthy cells are damaged by the drugs, intensive efforts are made to develop cancer-targeted techniques. The most promising approach is genetically modified CAR-T cell therapy. The high central memory T cell (Tcm) and stem cell-like memory T cell (Tscm) ratios in the CAR-T cell population increase the effectiveness of immunotherapy. Therefore, it is important to increase the populations of CAR-expressing Tcm and Tscm cells to ensure that CAR-T cells remain long-term and have cytotoxic (anti-tumor) efficacy. In this study, we aimed to improve CAR-T cell therapy’s time-dependent efficacy and stability, increasing the survival time and reducing the probability of cancer cell growth. To increase the subpopulation of Tcm and Tscm in CAR-T cells, we investigated to produce a long-term stable and cytotoxic efficient CAR-T cell by modifications in the cell activation-dependent production method using Phytohemagglutinin. Phytohemagglutinin (PHA), a lectin that binds to the membranes of T cells and increases metabolic activity and cell division, is studied to increase the Tcm and Tscm population. Although it is known that PHA significantly increases Tcm cells, B-lymphocyte antigen CD19 specific CAR-T cell expansion, its anti-cancer and memory capacity has not yet been tested compared to aCD3/aCD28. Two different types of CAR (aCD19 scFv CD8- (CD28 or 41BB)-CD3z-EGFRt) expressing T cells were generated and their immunogenic phenotype, exhausted phenotype, Tcm-Tscm populations, and cytotoxic activities were determined in this study. The proportion of T cell memory phenotype in the CAR-T cell populations generated by PHA was observed to be higher than that of aCD3/aCD28-amplified CAR-T cells with similar cytotoxic (anti-tumor) and higher proliferation capacity. Here, we show that PHA provides long-term and efficient CAR-T cell production, suggesting a potential alternative to aCD3/aCD28-amplified CAR-T cells.

## INTRODUCTION

The development of genetic modification techniques has opened a new era in cancer treatments that were limited to conventional treatments such as chemotherapy and monoclonal antibodies. The main reason is that the host immune cells cannot be sufficiently activated by cancer cells to exert a cytotoxic function due to low anti-tumor activity or lack of effector T cells (Frey, 2015). The most promising of these methods is genetically modified CAR-T cell therapy. The first clinical studies using transgenic CAR-T cells were in patients with hematological cancers, including B-cell acute lymphoblastic leukemia (ALL), Non-Hodgkin’s Lymphoma (NHL), chronic lymphocytic leukemia (CLL), and Multiple Myeloma (MM). This demonstrated improved response rates ranging from 50% to 85%, significant disease-free and overall survival (Finney et al., 1998; Sadelain et al., 2013). The US Food and Drug Administration (FDA), European Union, and Canada have approved Kymriah (Tisagenlecleucel) by Novartis for pediatric and young adult patients with ALL; also approved Yescarta (Axicabtagene ciloleucel) by Kyte Gilead for the treatment of adults with relapsed or refractory large B-cell lymphoma.

Chimeric antigen receptors are engineered receptors that typically contain the antigen binding site of a monoclonal antibody (mAb), T cell receptor transmembrane domain, and an intracellular signaling domain of the CD3ζ chain. Following initially disappointing results with the first generation of CAR-T cells, the cytotoxicity, expansion, and persistence of CAR-T cells in clinical studies have been improved in subsequent generations of CARs by including one or more intracellular domains of co-stimulatory molecules, such as CD28 or 4-1BB (Sadelain et al., 2015; Chang et al., 2017; Zhang et al., 2017). Trials with CARs containing CD28 or 4-1BB domains have shown similar initial response rates in patients with ALL (Lee et al., 2015; Maude et al., 2014; Sadelain et al., 2013). Long-term expression and effector function of CAR (CD28)-T cells were observed *in vivo* studies at 30 days. In contrast, the prolonged expressions of the effector function of CAR (4-1BB)-T cells have been reported to persist for up to 4 years in patients (Porter et al., 2015). In addition, the inclusion of 4-1BB signaling domains in some CARs has been shown to attenuate immunodeficiency (exhaustion) in CAR-T cells (Long et al., 2015). Endogenous 4-1BB signaling has been reported long-term survival benefits for T cells (Sabbagh et al., 2008).

The prolonged proliferation and durability that central memory T cells (Tcm) and stem cell-like memory T cells (Tscm) induce following T-cell therapy indicate crucial preconditions for the effectiveness of the treatment (Blaeschke et al., 2018). Literature studies have shown that the high proportions of Central memory T cells and stem cell-like memory T cells in the CAR-T cell population provide the necessary prerequisites (continuous proliferation and long-term persistence) for the effectiveness of immunotherapy (Blaeschke et al., 2016, 2018). Tcm and Tscm cell populations were found to be higher in the 4-1BB CAR group compared to the CD28 CAR group. In contrast, an increase in the T cell group with a high effector memory phenotype (Effector memory T cell, Tem) was observed in CD28 CAR cultures (Martinez-Forero et al., 2013; Sabbagh et al., 2008). A crucial phase of the adaptive immune response is T-cell activation. In addition to producing cytotoxic T-cell responses, activation is necessary for efficient CD4 T-cell responses (Soskic et al., 2014). Activated T cells may proliferate, differentiate, secrete cytokines, kill target cells, and perform other effector tasks (Lever et al., 2014). Clinically applicable conventional T-cell stimulation *in vitro* requires multivalent anti-CD3 and anti-CD28 antibodies (Pène et al., 2003).

As a good alternative, Phytohemagglutinin (PHA) binds to the TCR/CD3 complex, mimicking all intracellular activation events triggered by anti-CD3 antibodies (Faguet, 1977). PHA is a lectin derived from red kidney beans that bind to the membranes of T-cells and stimulates metabolic activity and cell division (Movafagh et al., 2011). PHA is a common polyclonal stimulant used in retroviral transduction protocols (Duarte et al., 2002). Human CD8+ T cells are known to be difficult to culture for extended periods (> 3 weeks) in culture. The optimal viability of CD8+ T cells can be maintained for up to 21 days with IL-15 in culture, but studies have shown that CD8+ T lymphocytes can be cultured for a long time (up to 90 days) by combining IL-15 and PHA (Isabel Da Cunha, M., and Terra, M., 2012). PHA induced a marked increase in the CD8+ Tcm population and CD8+ TN cells, more importantly, the CD8+ CCR7 Tem and TEf subsets exhibited the effector phenotype including cytotoxic capacity and performance expression (Supimon et al., 2021). Despite inducing cell proliferation, PHA also promotes cell apoptosis due to FasL interactions (Kabelitz and Janssen, 1997), which can be inhibited by IL-15 (Bulfone-Paus et al., 1997). Thus, the combination of PHA and IL-15 signals creates a favorable environment for the growth and expansion of CD8+ T cells.

In this research, Tcm and Tscm population ratios and cytotoxic activity were assessed in the PHA or aCD3/aCD28-activated CAR-T cells with two different CAR(aCD19 scFv-CD8-(CD28 and/or 41BB)-CD3z-EGFRt) constructs. The efficiency capacity was determined in CAR-T cell production and proliferation by adding IL-2, IL-7, IL-15 cytokines, and PHA in the medium. It is shown that cultured CAR-T cells with PHA give Tcm and Tscm profiles to these cells so that they can remain long-term and cytotoxic *in vitro* cancer models. With this research, the long-term and effective CAR-T cell production method will be tested in animal experiments.

## MATERIALS & METHODS

### Synthesis of CAR Construct and Lentivirus Production

The lentiviral vector encoding the CD19-CD28z specific CAR (CD19-CD8-CD28-CD3z) was designed and synthesized by Creative Biolabs. The lentiviral vector encoding CD19 specific CAR (CD19-CD8-4-1BB-CD3z) was designed and synthesized by GenScript. The envelope pCMV-VSV-G plasmid (from Bob Weinberg (Addgene #8454; http://n2t.net/addgene:8454; RRID: Addgene_8454)) and psPAX2 plasmid (from Didier Trono (Addgene #12260); http://n2t.net/addgene:12260; RRID: Addgene_12260)) required for lentivirus production were obtained from Addgene. Plasmids were amplified by integrating CAR-encoding plasmids to be used for genetic modification and plasmids (VSVG and psPAX2) used to produce lentivirus packaging proteins into *E. coli* DH5α (NEB C2987H) strain. Plasmid DNA was produced by using CompactPrep Plasmid Maxi Kit (QIAGEN, Cat No: 12863) and Zymopure Plasmid Maxiprep Kit (Zymopure, Cat: D4202). DNA concentration was measured in ng/µl with Microplate ELISA Reader (FLUOstar Omega) and it was evaluated whether the purity value was between 1.8<A260/A280<2.0. For the control of the isolated plasmids, DNA samples loaded on the 1% agarose gel prepared in 1X TAE buffer solution using BIO-RAD gel electrophoresis system were run at 90 V and 60 minutes. Plasmid DNA samples with <1% of bacterial DNA contamination were used for lentivirus production. Isolated envelope, packaging, and CAR plasmids were treated with FuGENE (Promega, E2311) transfection reagent and then lentivirus production was performed using host cells HEK293T [10% FBS and 1% penicillin/streptomycin, Gibco DMEM HG medium with L-Glutamine]. Packaged recombinant lentiviruses were collected 72 hours after transfection from the supernatant of HEK293T cell cultures. Produced CAR lentiviruses were concentrated with the Lenti-X Concentrator (Takara Bio, 631232) to increase virus concentration (20X-100X). Viruses were stored at −80° C (Tastan et al., 2020).

### Lentivirus Titration

The Jurkat cell line was suspended as 10,000 cells per well in 100 µl of RPMI with glutamine HEPES including 10% FBS, 1% pen/strep, 1% non-essential amino acids, 1% sodium pyruvate, and 1% vitamins. Jurkat cells in 100 µl of medium were plated in U bottom 96-well plates from A to H. The wells were adjusted to have 10 µl, 3 µl, 1 µl, 0.3 µl, 0.1 µl, 0.03 µl, and 0.01 µl of the 100x-concentrated CAR-LV solutions in each 50 µl of the medium, respectively, and then 50 µl of virus dilution from each concentration was transferred to Jurkat cultured wells, the total volume was adjusted to 150 µl, and cells were incubated for 72 hours. EGFRt expression was determined using Cytoflex Flow Cytometer (Beckman Coulter, B5-R3-V0) with an anti-EGFR-FITC antibody (R&D Systems, FAB10951G).

### T Cell Transduction and CAR-T Culture Conditions

The research was approved by the Acibadem University and Acibadem Health Institutions Medical Research Ethics Committee (ATADEK-2019-17/31). Healthy adult blood samples were obtained and peripheral blood mononuclear cell (PBMC) isolation was performed at Uskudar University, Transgenic Cell Technologies and Epigenetics Application and Research Center (TRGENMER). Blood samples from three healthy human donors are combined with Ficoll (Paque PREMIUM 1.073, 17-5446-52). PBMCs are isolated by the density-gradient centrifugation method. Total CD3+ T cells were isolated using anti-CD3 microspheres (Miltenyi Biotech, 130-050-101). Initial T cell activation was performed with anti-CD3/anti-CD28 microbeads (T Cell TransAct, Miltenyi Biotech, 130-111-160) and 10µg/ml Phytohemagglutinin-M (PHA-M) (Roche 11082132001, 20 mg). Lentivirus transduction procedure, Vectofusin 1 (10µg/ml) (Miltenyi Biotech, 130-111-163), and BX-795 hydrochloride (6µM) (Millipore Sigma, SML0694-5MG) were inoculated with 2-3 MOI lentiviruses. Cells were cultured in T cell medium (50 IU/ml IL-2, 10 ng/ml IL-7, 5 ng/ml IL-15, 10% Fetal Bovine Serum and 1% pen/strep) for 14 days. CAR expression levels were determined by Cytoflex Flow Cytometer analysis using an anti-EGFR-FITC antibody. The second activation and expansion (re-stimulation) study on day 14 was performed using PHA and aCD3/aCD28. Re-transduction assay was performed using Vectofusin-1 with 5-10 MOI lentiviruses 24 hours after the second activation (Tastan et al., 2020).

### Analysis of T Cell Sub-Populations (Tn-Tcm-Tem-Tef) and Exhaustion Profile

T cell sub-population profiling studies on days 14 and 21 were analyzed using CD3-PC7 (Beckman Coulter, 6607100), CD4-APC-A700 (Beckman Coulter, B10824), CD8-PC5.5 (Beckman Coulter, B21205), EGFR-A488 (R&D Systems, FAB10951G), CD45RA-ECD (Beckman Coulter, IM2711U), CD45RO-PE (Miltenyi Biotech,130-113-559), CD27-APC-A750 (Beckman Coulter, B12701) and CD62L-APC (Miltenyi Biotech, 130-113-617) antibodies by Cytoflex Flow Cytometer analysis. Exhaustion profiling studies on days 14 and 21 were analyzed using CD3-PC7 (Beckman Coulter, 6607100), CD4-APC-A700 (Beckman Coulter, B10824), CD8-PC5.5 (Beckman Coulter, B21205), EGFR-A488 (R&D Systems, FAB10951G), CD279 (PD1)-PE (Miltenyi Biotech, 130-117-384), CD366 (TIM3) - APC (Invitrogen, 17-3109-42) and CD223 (LAG-3) - APC-eFluor 780 (Invitrogen, 47-2239-42) antibodies by Cytoflex Flow Cytometer analysis.

### *In Vitro* Anti-Tumor Cytotoxicity and Efficacy Assay

*In vitro* studies with CAR-T cells (CD19-CD28z and CD19-BBz) were performed for the assessment of efficacy and cytotoxic capacity. For that purpose, anti-CD19-expressing CAR-T cells and CD19-expressing RAJI cells were cultured for 24 h, 7 days, and 14 days (CAR-T: RAJI; 1:1, 5:1, 10:1). Anti-Cancer profiling studies on days 14 and 21 were analyzed using CD3-PC7 (Beckman Coulter, 6607100), CD4-APC-A700 (Beckman Coulter, B10824), CD8-PC5.5 (Beckman Coulter, B21205), EGFR-A488 (R&D Systems, FAB10951G), CD19-ECD (Beckman Coulter, A07770), CD25-APC (Miltenyi Biotech, 130-113-284), CD107a (LAMP-1) -PE (Miltenyi Biotech, 130-111-621) by Cytoflex Flow Cytometer analysis. In the co-culture experiments, the death of CD19+ RAJI cells after 48 hours and the CD25 activation (IL2RA, IL-2 receptor alpha chain) and CD107a (marker for degranulation of lymphocytes) cytotoxic de-granulation biomarkers of CAR-T cells and control T cells in CD3+ T cells were analyzed by flow cytometry. Survival analysis of CAR-T cells was performed to control the cell viability of RAJI cells and CAR-T cells. Trypan blue (Biological Industries, #03-102-1B) was applied to identify and count surviving cells. Cell counting and viability analysis were performed with the BIO-RAD TC20 Automated Cell Counter.

### Statistics Analysis

Two-tailed Homoscedastic T-tests were performed using SPSS software. Outliers were not excluded in any of the statistical tests and each data point represents an independent measurement. Bar plots report the mean and standard deviation or the standard deviation of the mean. The threshold of significance for all tests was set at p*<0.05.

## RESULTS

### Isolation of CD3+ T Cells

For the isolation of CD4+ and CD8+ T cells from PBMC, CD3+ T cell isolation was performed by using anti-CD3 microspheres (Miltenyi Biotech, 130-050-101) instead of anti-CD4 and anti-CD8 microspheres. Based on the Tcm-Tscm ratios we obtained in this research, we aimed to develop the CAR-T cell production protocol with CD3 isolation. We have successfully achieved >99% CD3 rate in CD3+ T cell isolations with 3 different donors **(Figure-1)**.

**Figure 1:**
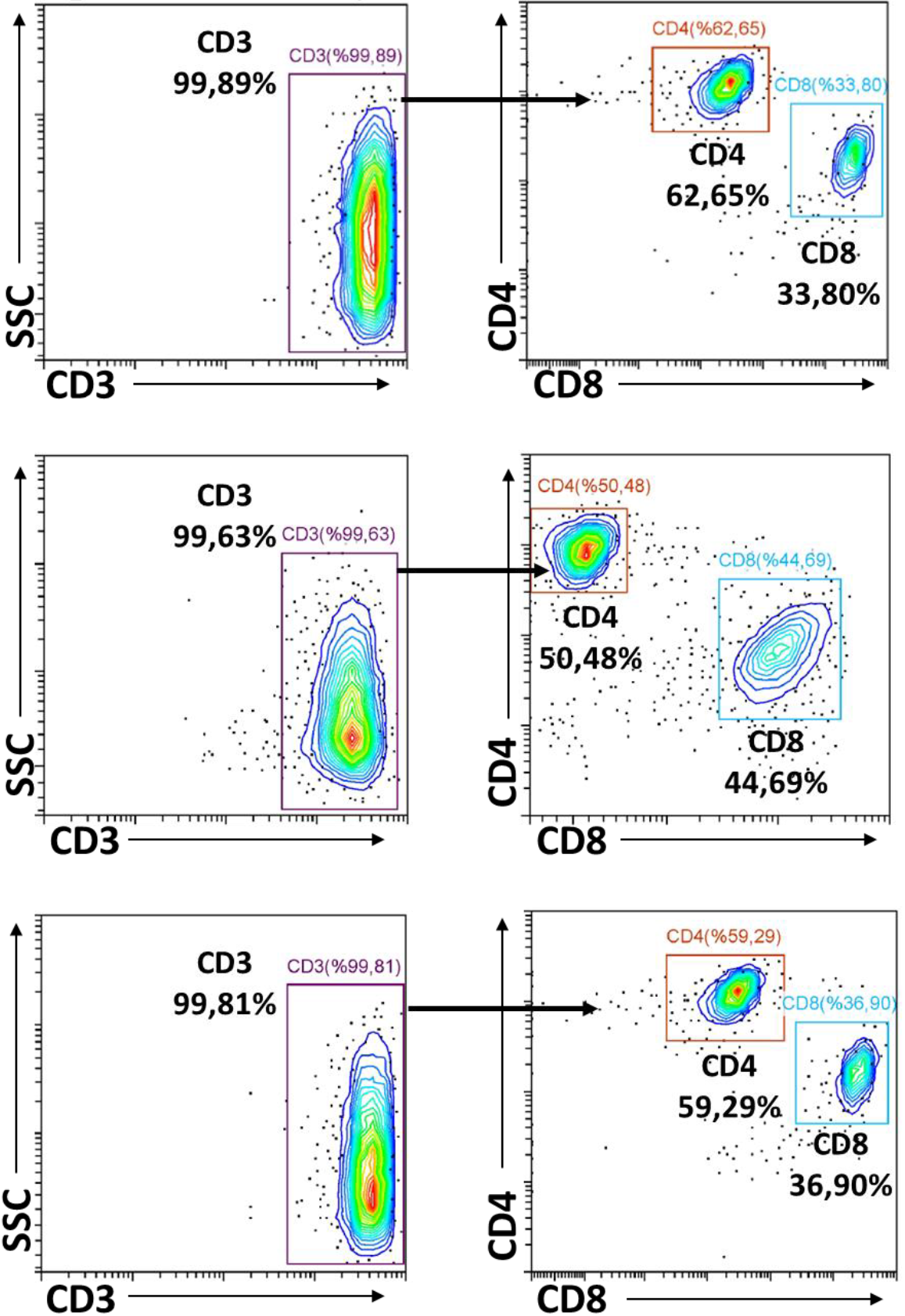
The Flow cytometry Plots of CD3+ T cell isolation from 3 different donors.

### *In Vitro* Assessment of CAR-T Cell Proliferation, Differentiation, and Anti-Cancer Capacity

In our research, the activation of T cells was activated was performed by PHA (10µg/ml) or the use of anti-CD3/anti-CD28. It was questioned whether PHA increases the Tcm and Tscm population considering anti-CD3/anti-CD28 microbeads. CAR-T cell production, activity, and sub-population profiling following T cell activation with PHA were determined **(Figure 2)**.

**Figure 2:**
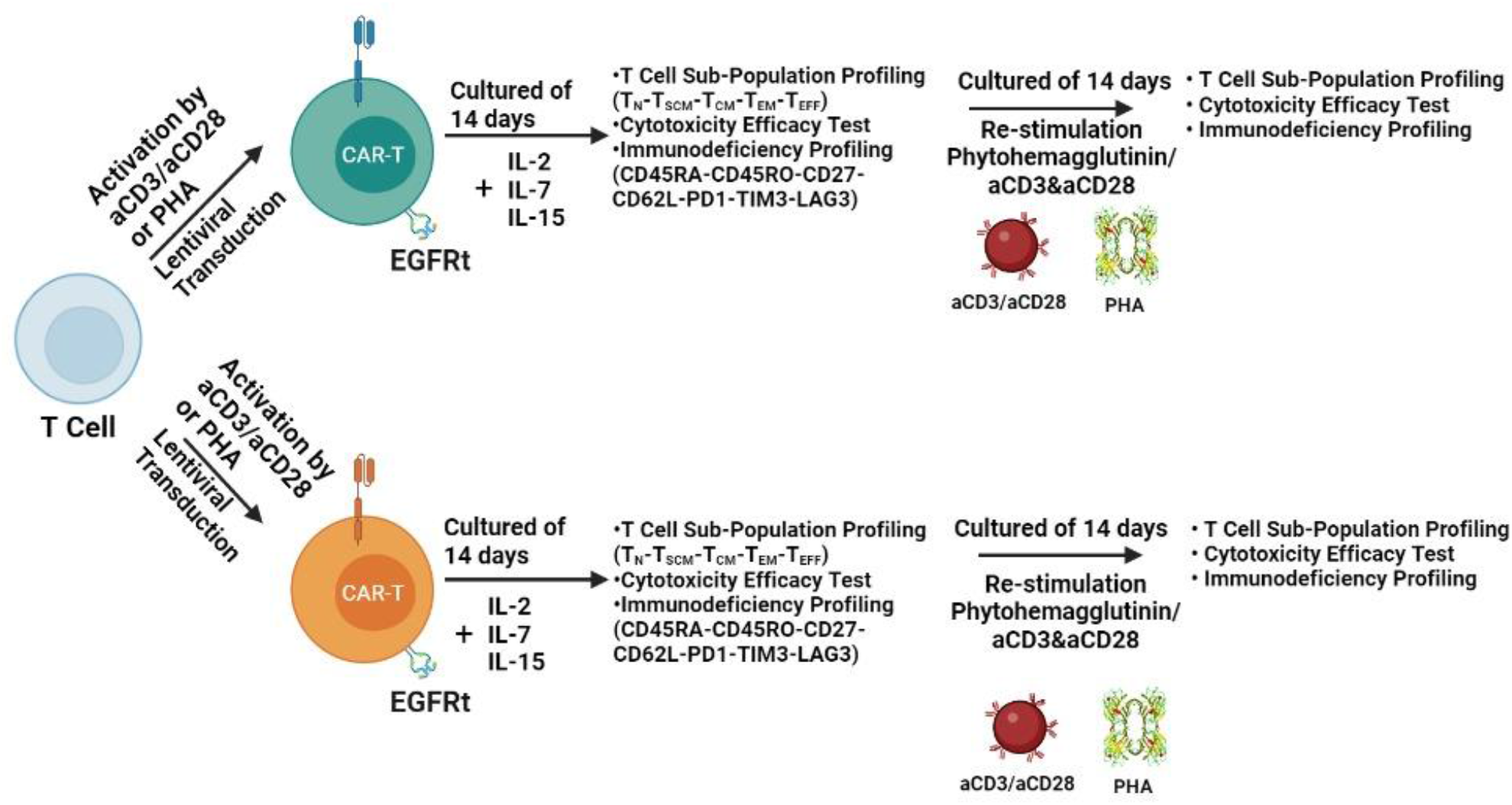
The experimental setup aiming to determine Tcm and Tscm sub-population, proliferation, and cytotoxicity of CAR-T cells with 4-1BB or CD28-based CAR construct upon activation of T cells with Phytohemagglutinin/ aCD3&aCD28.

CAR expressions of CAR-T cells obtained by culturing CD3+T cells transduced with CAR19BB and CAR1928 lentiviruses with Anti-EGFR-A488 antibody were analyzed by flow cytometry. As a result of the analysis, we evaluated CAR expression rate >20% in CAR (4-1BB)-T and CAR (CD28)-T cells (**Figure 3**). CAR-T cells activated with anti-CD3/anti-CD28 and PHA and transduced with CAR19BB had an expression rate of 22.31% and 22.06%, respectively. CAR-T cells activated with anti-CD3/anti-CD28 and PHA and transduced with CAR1928 had an expression rate of 19.53% and 12.70%, respectively.

**Figure 3:**
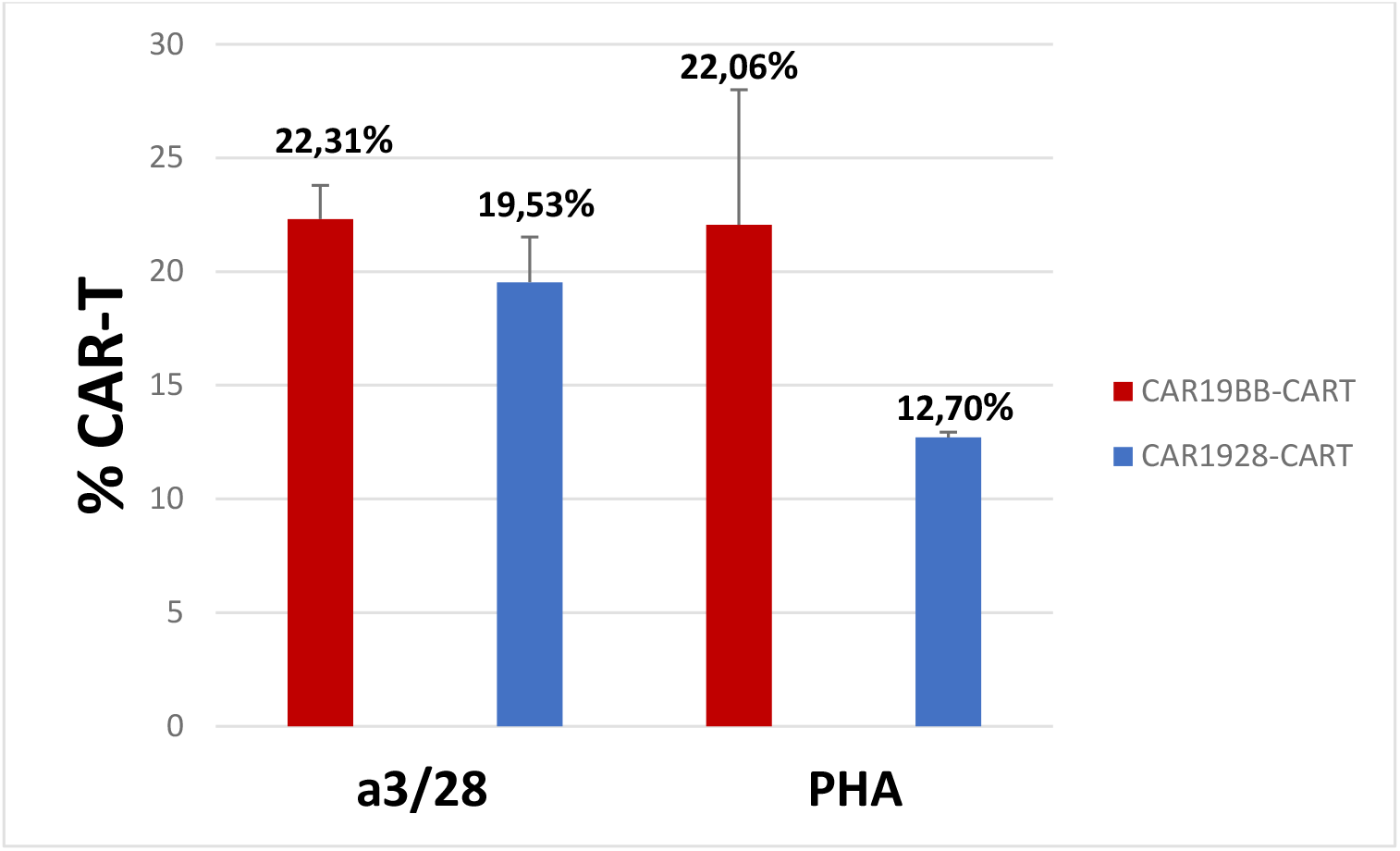
The bar graph showing CAR+ T cell frequency with CAR19BB or CAR1928 constructs after activation with PHA or anti-CD3/anti-CD28 by expression of EGFRt.

With the use of CAR19BB lentivirus, we have completed the production of CAR-T cells after activation with both anti-CD3/anti-CD28 and PHA. Apart from this, although we approached more than 20% CAR expression with anti-CD3/anti-CD28 activated CAR1928 T cells, we could not achieve this with PHA (±20% standard deviation). Due to this situation, we have decided to increase our MOI ratios from 2-3 MOIs to 5-10 MOIs to provide more than 20% CAR expression in cells in a second transduction process.

The second activation and expansion (re-stimulation) study on day 14 was performed using PHA and aCD3/aCD28. PHA/aCD3/aCD28-activated CD3+T cells were reactivated with PHA/aCD3/aCD28. The second transduction was performed 24 hours after the re-activation. Thus, we tested whether the use of 5-10MOI of CAR lentivirus would increase our expression by second transduction. When the CAR expressions in T cells were assessed, it was observed that the PHA-activated T cells expressed 72.5± 12.3% CAR while anti-CD3/anti-C28 activated T cells expressed 58.7± 4.1% (**Figure 4**). Thus, when we activated it with PHA, we ensured the production of CAR-T cells effectively.

**Figure 4:**
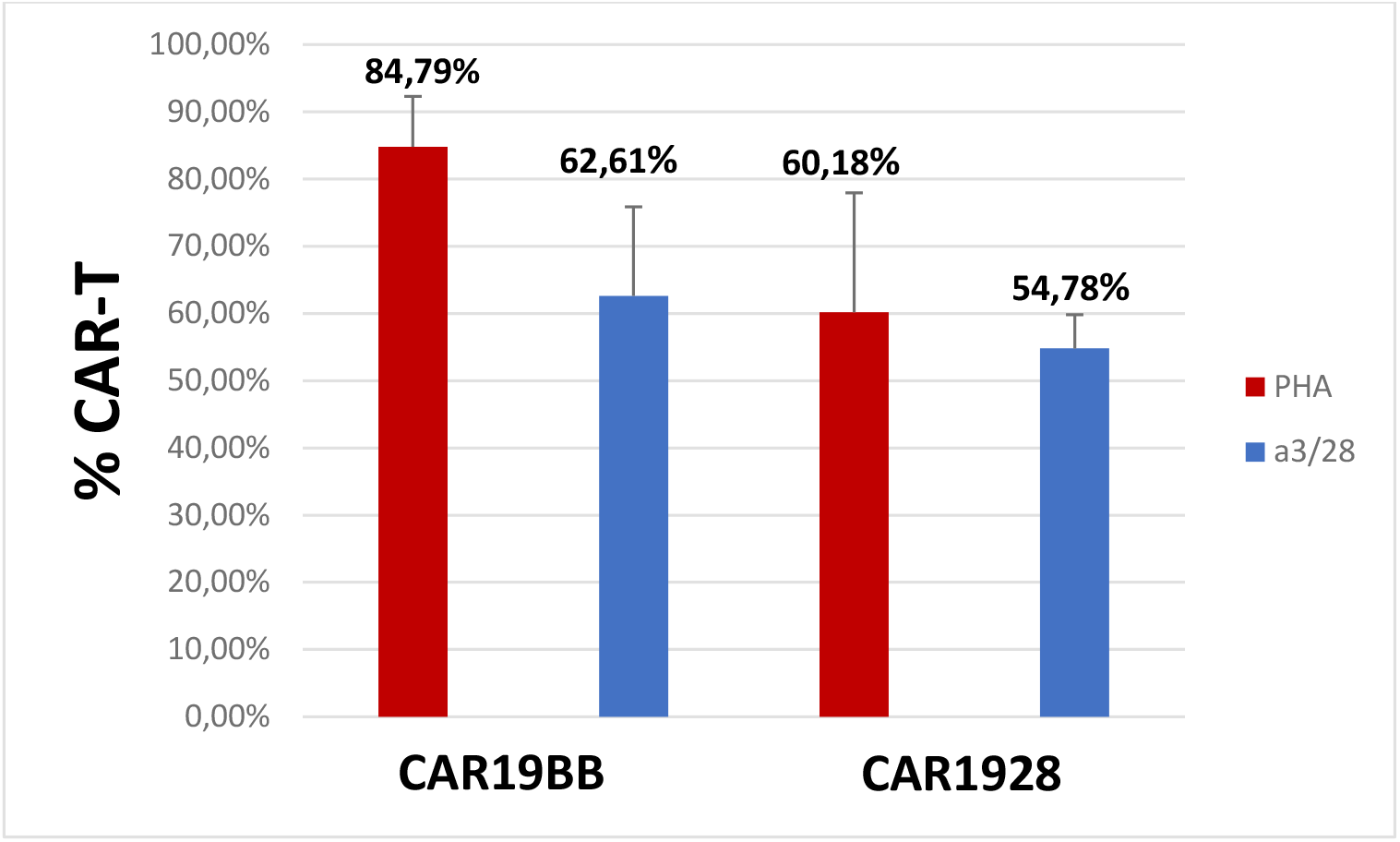
The bar graph showing CAR+ T cell frequency with CAR19BB or CAR1928 constructs after second activation and re-transduction with PHA or anti-CD3/anti-CD28 by expression of EGFRt.

In the next experiment, we investigated the proliferation capacity of CAR-T cells produced with PHA considering the anti-CD3/anti-CD28 activated CAR-T cells. The graph of the initial cell number of CAR-T cells activated with PHA and anti-CD3/anti-CD28 on day 0 and the division coefficient of cell proliferation on day 14 was determined in **Figure 5**. Here we show that the CAR-T cell proliferation capacity is significantly higher when the cells are stimulated with PHA (**Figure 5**).

**Figure 5:**
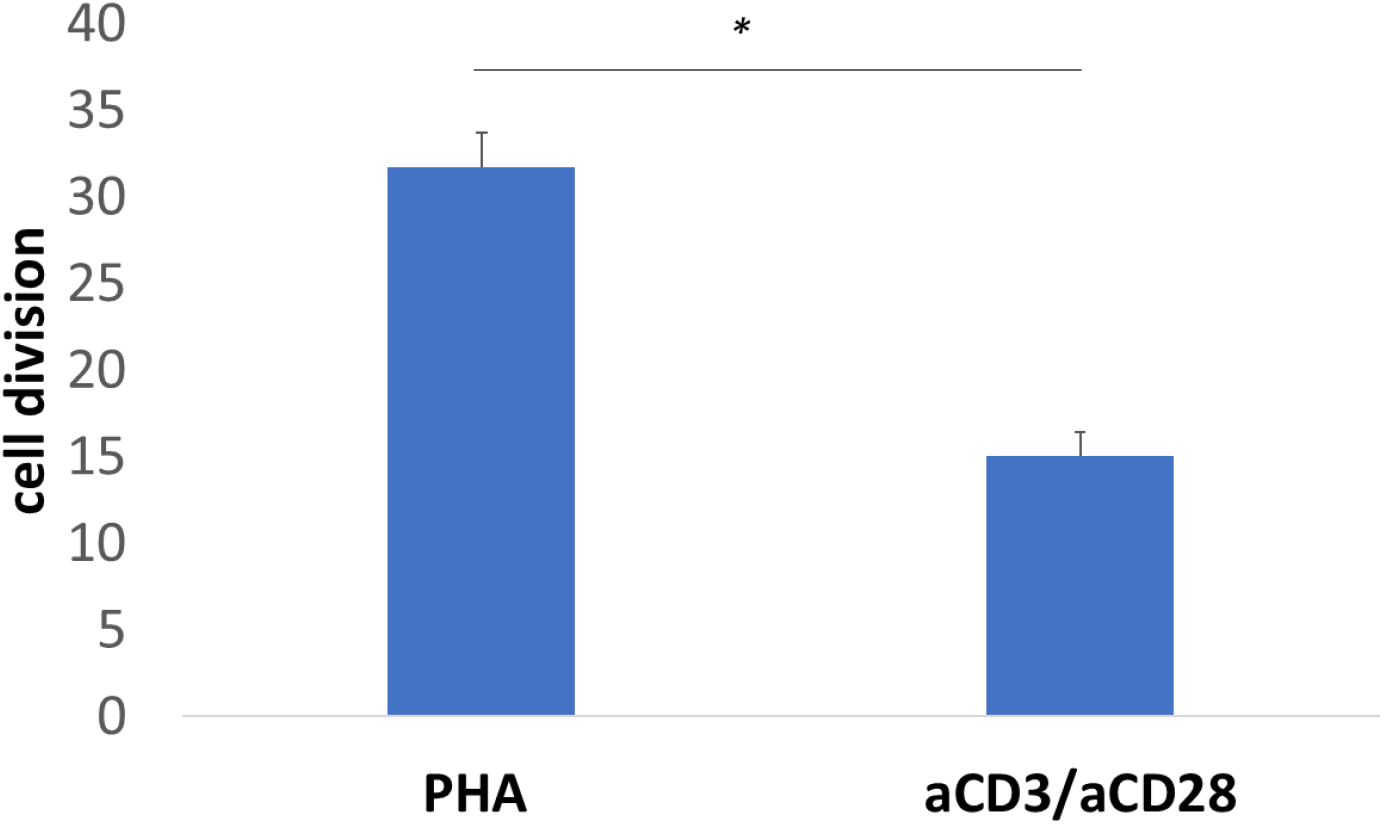
The bar graph showing the cell division capacities of the CAR-T cells stimulated either with PHA or aCD3/aCD28 on day 14.

### Immunoprofiling sub-populations of the activated T cell (Tn-Tcm-Tem-Tef)

The activated T cells differentiate to develop subtypes of effector and memory T cells. Next, we aimed to determine T cell sub-population frequencies upon activation with different stimulation reagents including PHA and aCD3/aCD28. To immunoprofile the activated T cells, we determine the Tcm-Tscm sub-populations using conventional immune biomarkers including Tn+Tscm (CD3^+^, CD45RA^+^, CD62L^+^), Tcm (CD3^+^, CD45RA^−^CD62L^+^), TemEarly (CD3^+^, CD45RO+, CD27+, CD45RA^−^, CD62L^−^) T cells, TemLate (CD3^+^, CD45RO+, CD27-, CD45RA^−^, CD62L^−^) and Temra (CD3^+^, CD45RA^+^CD62L^−^) (Ammirati et al., 2012; Larbi et al., 2014; Alvarez et al., 2016; Tuluc et al., 2017) (**Figure 6**).

**Figure 6:**
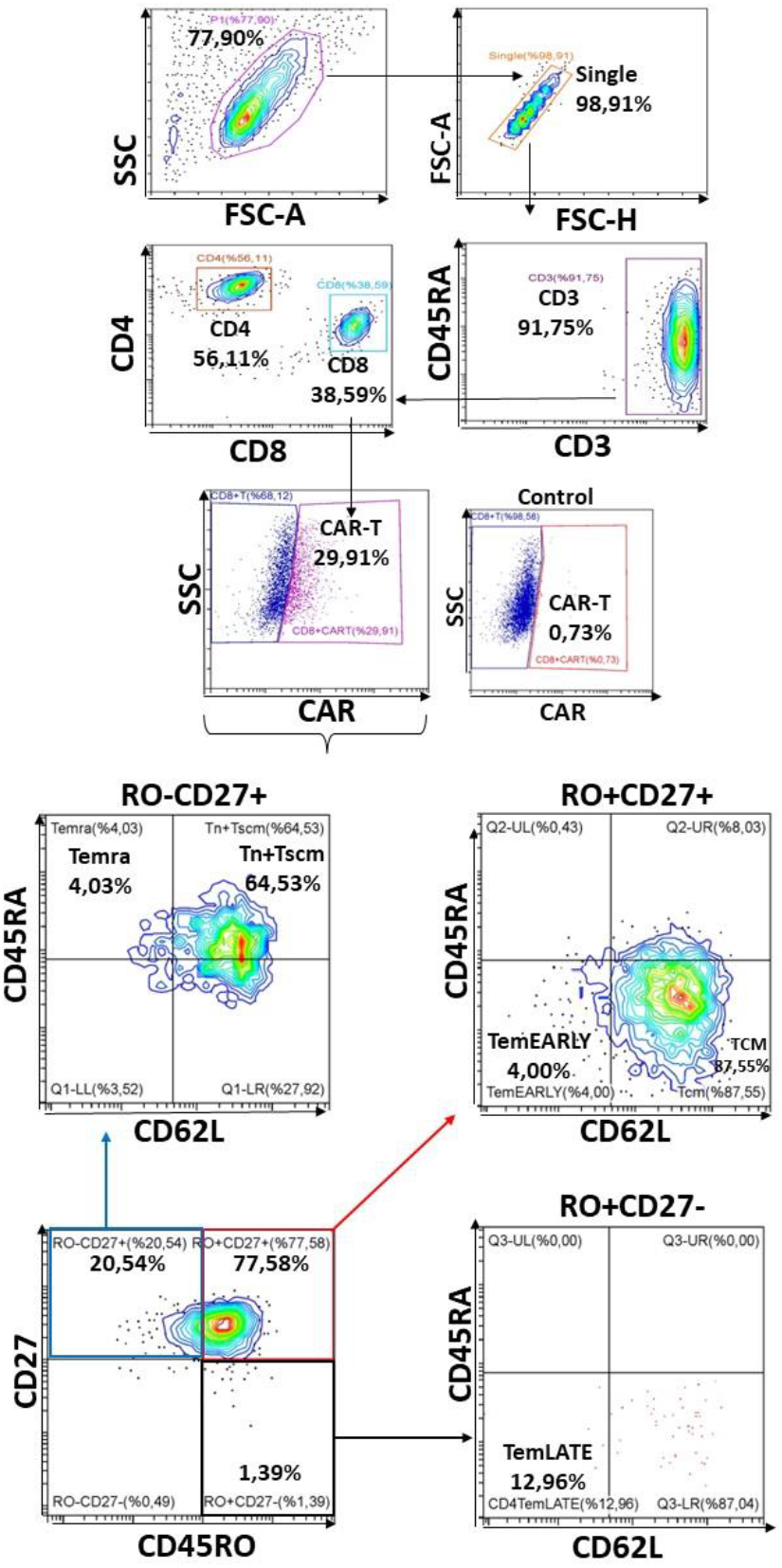
The flow cytometry plots showing the T cells sub-populations in CD4^+^ and CD8^+^ including **Tn+Tscm** (CD3^+^, CD45RA^−^, CD62L^+^), **Tcm** (CD3^+^, CD45RA^−^CD62L^+^), **TemEarly** (CD3^+^, CD45RO+, CD27+, CD45RA^−^, CD62L^−^) T cells, **TemLate** (CD3^+^, CD45RO+, CD27-, CD45RA^−^, CD62L^−^) and **Temra** (CD3^+^, CD45RA^+^CD62L^−^) in healthy donors.

In the examination of Tn+Tscm, Tcm, TemEARLY, TemLATE, and Temra sub-populations of total CD4+T cells among PHA-activated CD3+T cells, the Tn+Tscm ratio increases until day 14 **(Figure 7A and 7B)**. On day 14, the Tn+Tscm ratio increased. However, on day 21 after the second activation, the Tn+Tscm ratio decreased significantly **(Figure 7C)**. While the Tcm rate decreased until day 14, it increased statistically on day 21 after the second activation. Compared to day 7 and day 14, the rate of TemEARLY increases especially in the reactivation. While the Temra rate increases slightly until day 14, it decreases on day 21 **(Figure 7D)**.

**Figure 7:**
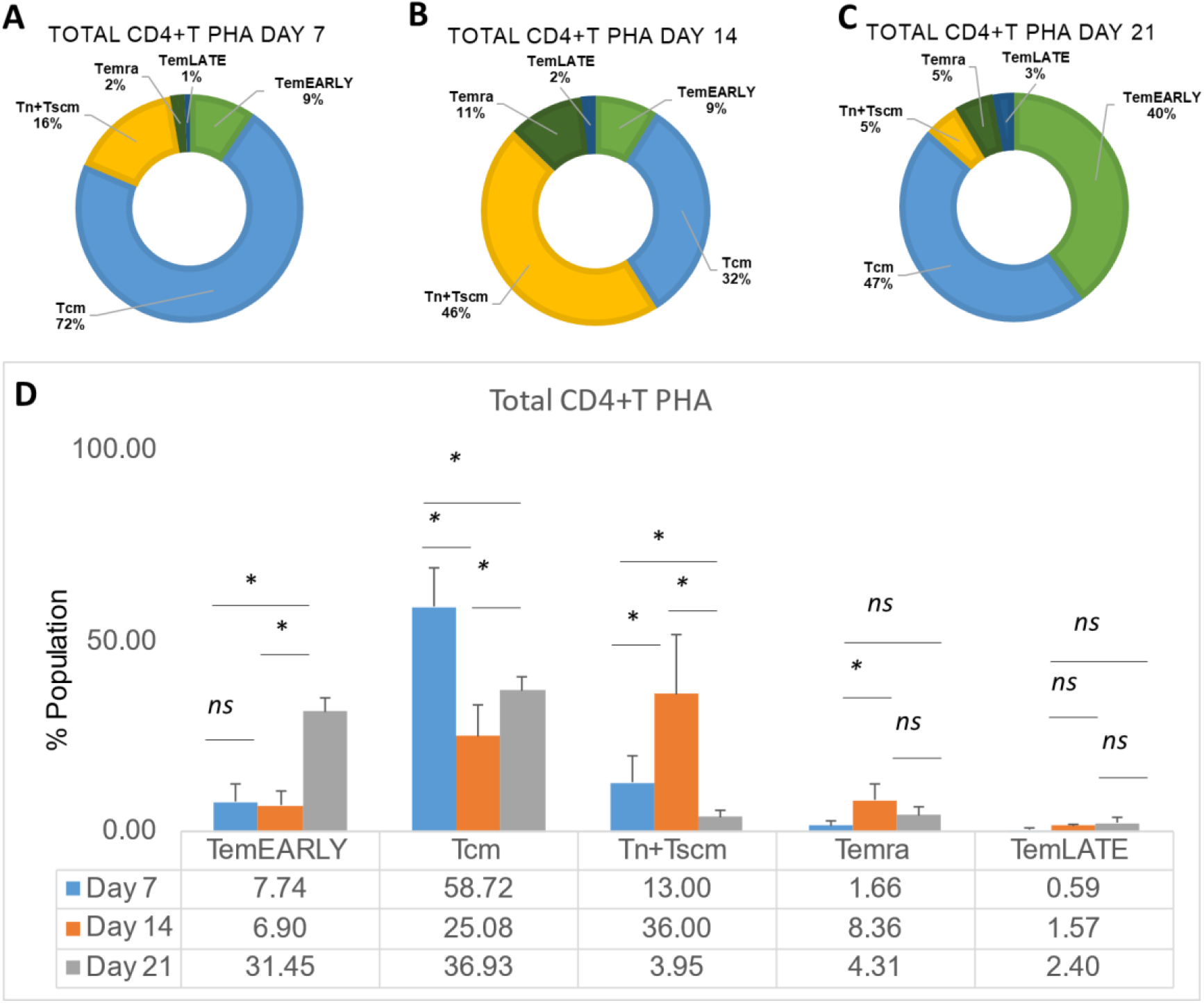
The pie charts showing the distribution of Tn+Tscm, Tcm, TemEARLY, TemLATE, and Temra sub-populations of total CD4+T cells in PHA-activated CD3+T cells on **A**. day 7, **B**. day 14, and **C**. day 21. **D**. The bar graph shows the quantitated distribution of the activated T cell sub-types.

In the examination of Tn+Tscm, Tcm, TemEARLY, TemLATE, and Temra sub-populations of total CD8+T cells among PHA-activated CD3+T cells, the Tn+Tscm ratio increases until day 14 after the first activation; while the Tcm rate decreases on day 14 compared to day 7 **(Figure 8A and 8B)**. TemEARLY increases after the reactivation **(Figure 8C)**. TemLATE also increases on day 21 compared to day 7 **(Figures 8A and 8C)**. An increase was observed in Temra until day 14 **(Figure 8D)**. Statistically, there is an increase in the rate of Temra in reactivation **(Figure 8D)**. Therefore, the increase in the exhausted PHA-activated cells seems to have differentiated after the reactivation.

**Figure 8:**
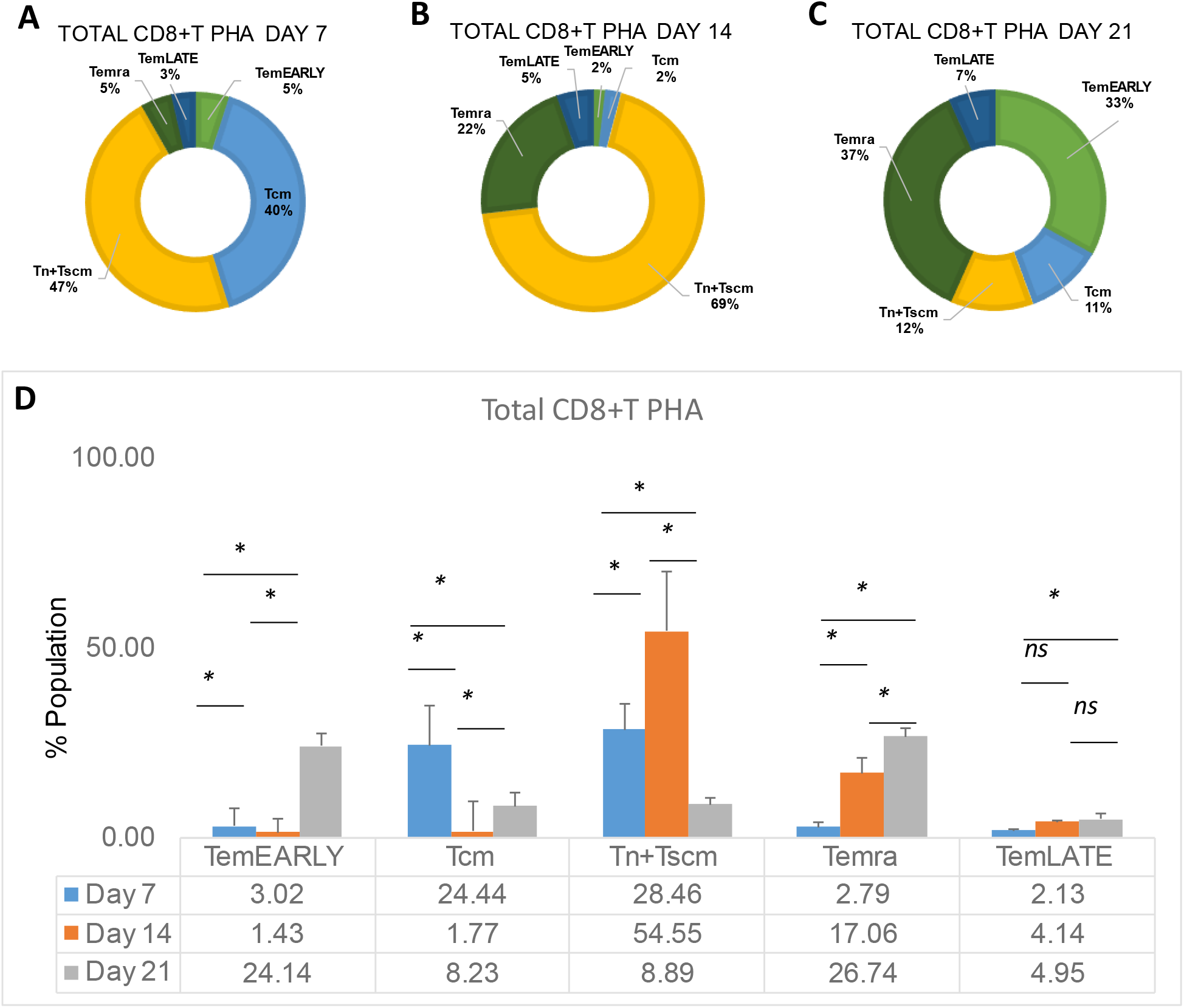
The pie charts showing the distribution of Tn+Tscm, Tcm, TemEARLY, TemLATE, and Temra sub-populations of total CD8+T cells in PHA-activated CD3+T cells on **A**. day 7, **B**. day 14, and **C**. day 21. **D**. The bar graph shows the quantitated distribution of the activated T cell sub-types.

To determine the population of Tn+Tscm, Tcm, TemEARLY, TemLATE, and Temra sub-populations of total CD4+T cells in anti-CD3/anti-CD28-activated CD3+T cells, the Tcm rate decreased at day 14 compared to day 7 **(Figures 9A and 9B)**. The Tcm rate increases on day 14 compared to day 14 **(Figure 9B)**. Although the Tn+Tscm ratio increased on day 14 after the first activation compared to day 7, it decreased on day 21 after the reactivation **(Figure 9A and 9C)**. While the rate of TemLATE increases from day 7 to day 14, it decreases on day 21 after the reactivation **(Figure 9D)**. The Temra rate also increased on day 14 and day 21 compared to day 7 **(Figure 9)**.

**Figure 9:**
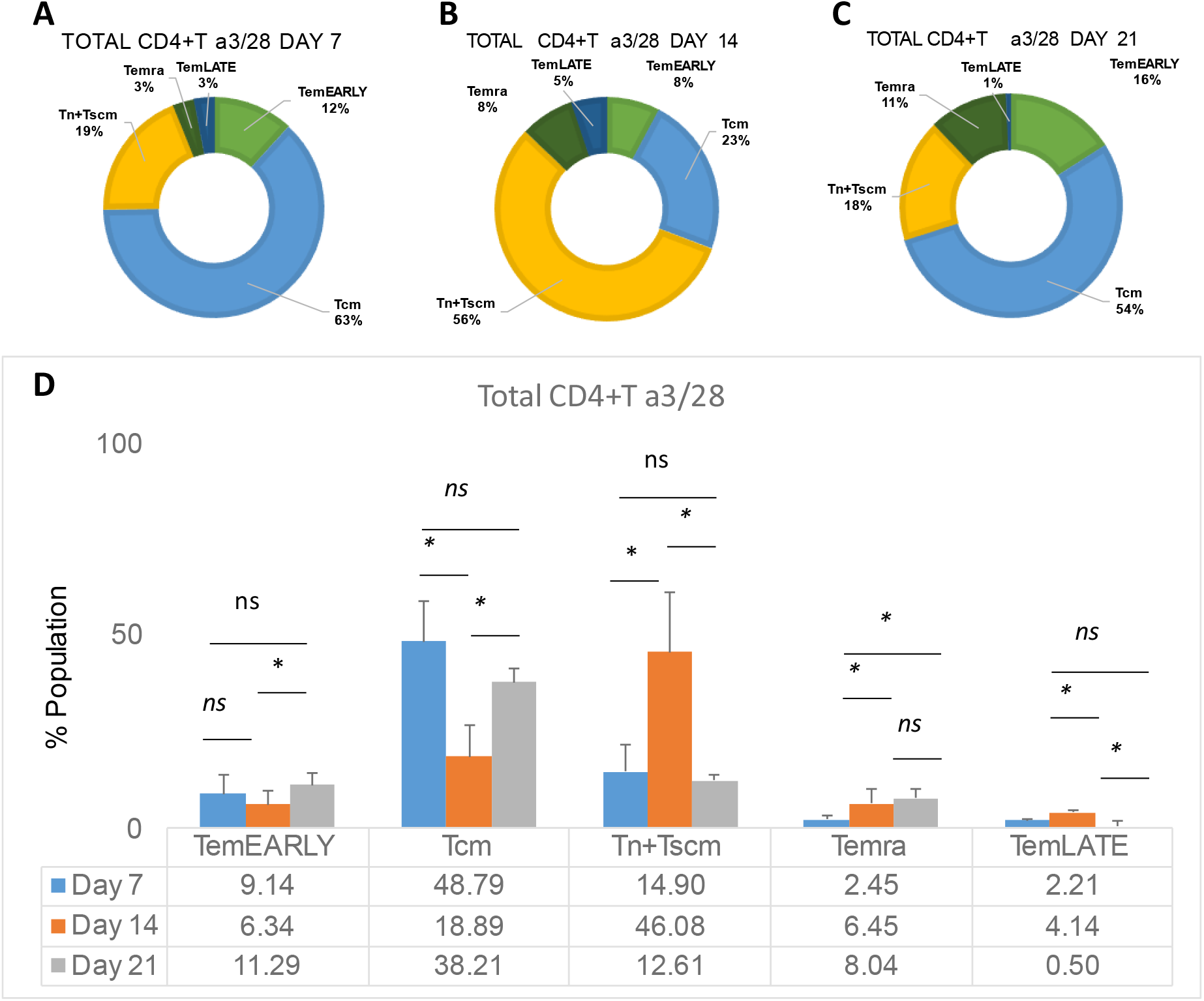
The pie charts showing the distribution of Tn+Tscm, Tcm, TemEARLY, TemLATE, and Temra sub-populations of total CD4+T cells in aCD3/aCD28-activated CD3+T cells on **A**. day 7, **B**. day 14, and **C**. day 21. **D**. The bar graph shows the quantitated distribution of the activated T cell sub-types.

To assess the frequency of Tn+Tscm, Tcm, TemEARLY, TemLATE, and Temra sub-populations of total CD8+T cells in CD3+T cells activated with anti-CD3/anti-CD28, the Tn+Tscm ratio increased on day 14 compared to day 7 **(Figures 10A and 10B)**. Then, on day 21, this rate decreases **(Figure 10C)**. The Tcm rate decreases on day 14 and day 21 days compared to day 7 **(Figure 10D)**. TemLATE and Temra rates increase statistically on day 14 and day 21 compared to day 7 **(Figure 10D)**.

**Figure 10:**
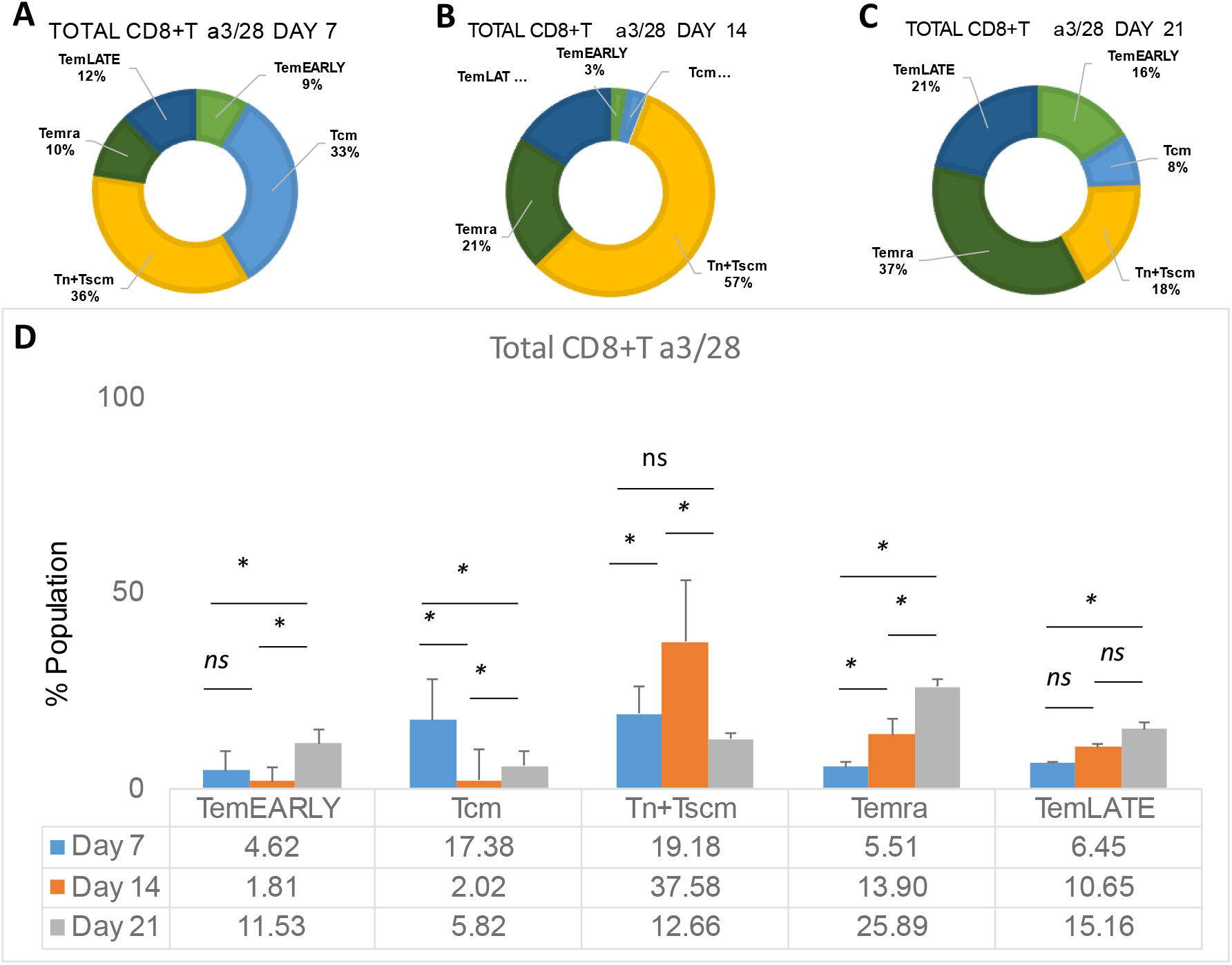
The pie charts showing the distribution of Tn+Tscm, Tcm, TemEARLY, TemLATE, and Temra sub-populations of total CD8+T cells in aCD3/aCD28-activated CD3+T cells on **A**. day 7, **B**. day 14, and **C**. day 21. **D**. The bar graph shows the quantitated distribution of the activated T cell sub-types.

PHA and anti-CD3/anti-CD28 activated CD3+T cells were compared. It was observed that there was no statistical difference for Tn+Tscm, Tcm, TemEARLY, and Temra except TemLATE in total CD4+T cells **(Figure 11)**. Therefore, the only difference was determined in the frequency of TemLATE with a decrease upon activation with PHA.

**Figure 11:**
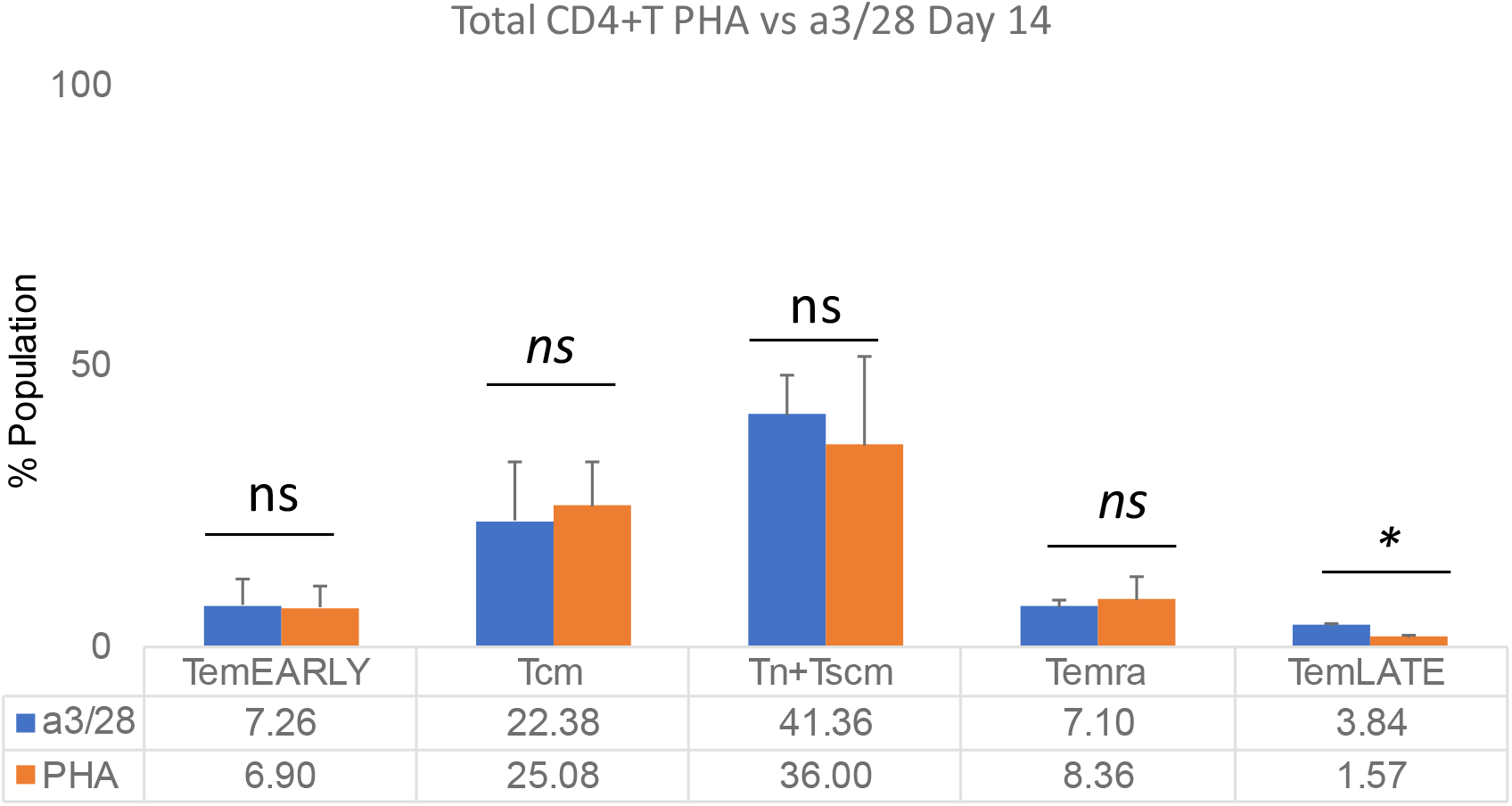
The bar graph showing the comparison of Tn+Tscm, Tcm, TemEARLY, TemLATE, and Temra sub-populations of total CD4+T cells either activated with anti-CD3/anti-CD28 or PHA at day 14.

When the effect of PHA and anti-CD3/anti-CD28 activation are compared, the Tn+Tscm ratio was statistically increased while the TemLATE frequency was impaired in the total PHA-induced CD8+T cells **(Figure 12)**. This data suggests that PHA-activated cells differentiate into early memory phenotypes rather than that with anti-CD3/anti-CD28 activation.

**Figure 12:**
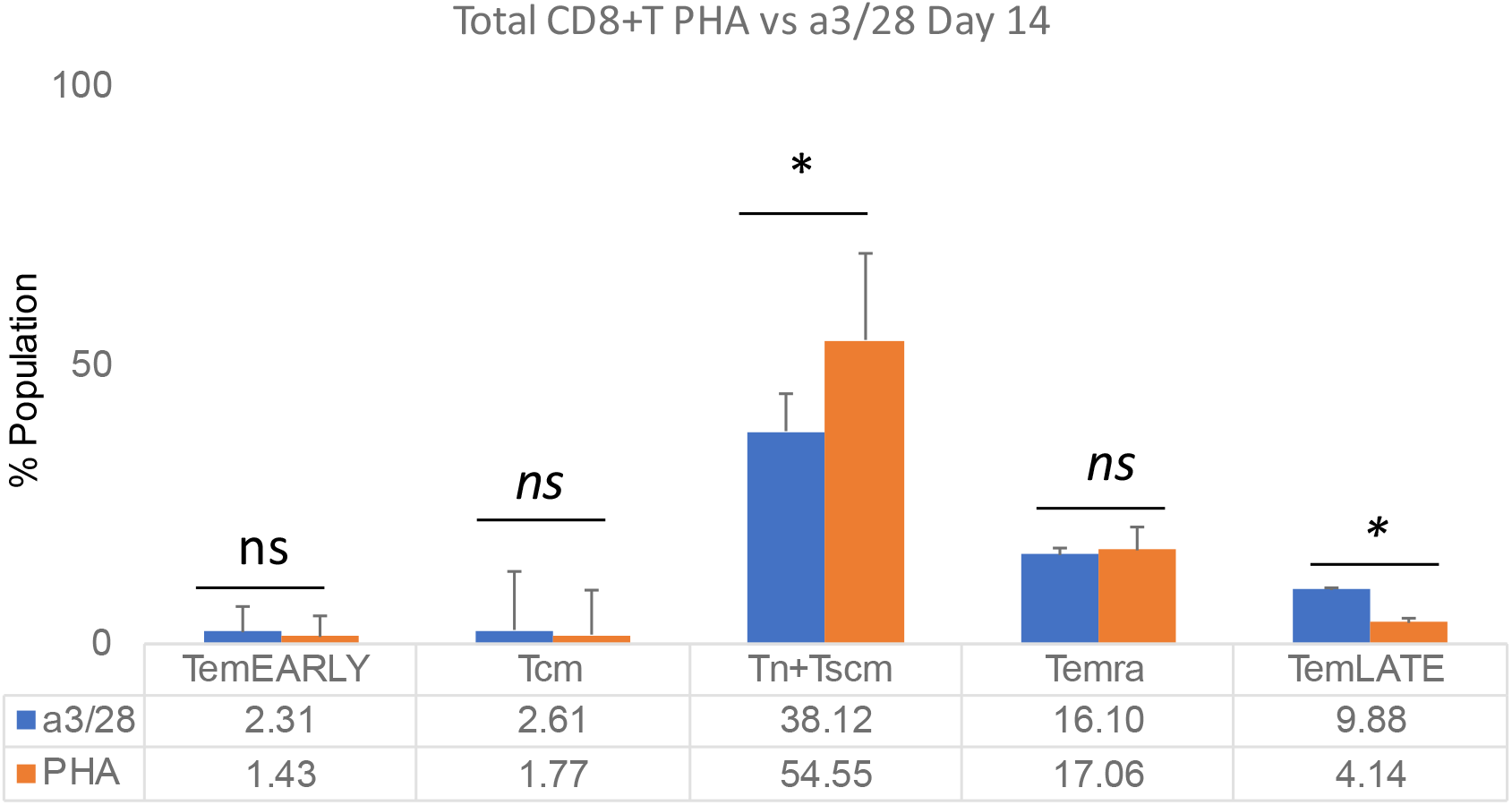
The bar graph showing the comparison of T-naive+Tscm, Tcm, TemEARLY, TemLATE and Temra sub-populations of total CD8+T cells either activated with anti-CD3/anti-CD28 or PHA on day 14.

Next, we aimed to investigate the change in distribution of CD4+ and CD8+T cell sub-types reactivated with PHA and anti-CD3/anti-CD28 on day 21. We determined that the Tn+Tscm population significantly decreased when the cells were activated with PHA **(Figure 13A)**. Although there was no change in Tcm population, PHA-activated TemEARLY and TemLATE populations were higher **(Figure 13A)**. When we determined the level of exhaustion markers on Temra, PHA-induced cells had lower levels rather than the anti-CD3/anti-CD28 cells **(Figure 13A)**. On the other hand, the ratio of Tn+Tscm and Tcm in total PHA-activated CD8+T cells was not significantly different from the anti-CD3/anti-CD28 activated cells (**Figure 13B**). However, the level of TemEARLY increased in the PHA-activated population. Also, the rate of TemLATE was significantly high in the anti-CD3/anti-CD28 activated population **(Figure 13B)**. These data suggest that PHA supports early memory phenotypes in the reactivated T cells.

**Figure 13:**
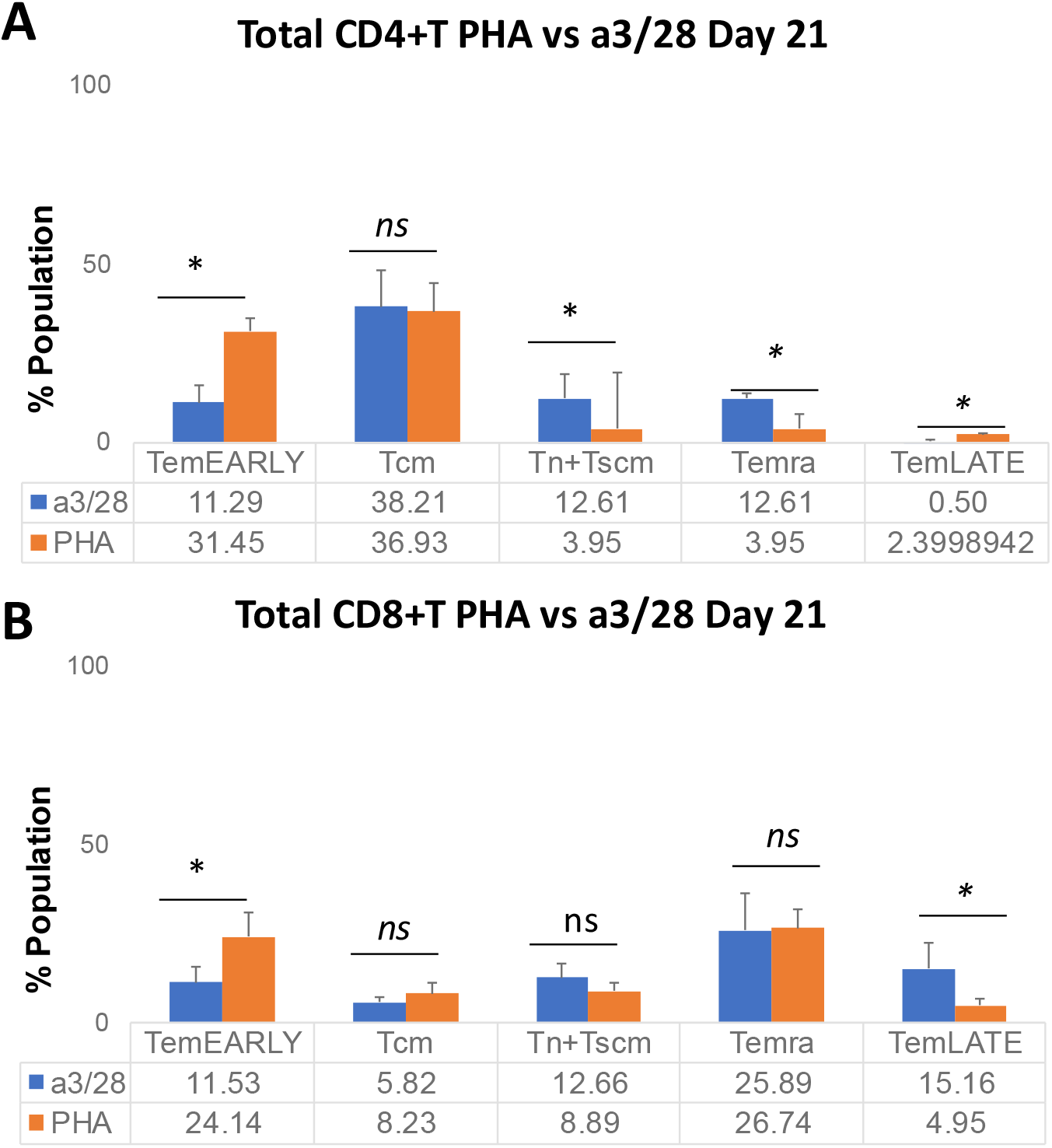
The bar graphs showing the Comparison of T-naive+Tscm, Tcm, TemEARLY, TemLATE, and Temra sub-populations of **A**. total CD4+T cells and **B**. total CD8+T cells activated either with anti-CD3/anti-CD28 or PHA on day 21.

Next, we wanted to determine the effect of CAR1928 or CAR19BB expression on T cells to differentiate into T cell subtypes. We determined subtype frequencies of the CD4+ or CD8+ T cells expression CAR constructs including CD28 or 41-BB costimulatory domain. On day 7 and day 14, there was no significant change in the subtype populations of the CAR T cells either with PHA and antiCD3/antiCD28 **(Figure 14A)**. On the other hand, upon restimulation, we determined that TemEARLY frequency was significantly high in the CAR19BB T cells activated with PHA **(Figure 14A)**. However, Tn+Tscm population level was impaired in the PHA-activated CAR19BB T cells considering antiCD3/antiCD28 activated cells **(Figure 14A)**. This data suggests that for the first activation and expansion, there is no significant effect of CAR construct expression for T cell differentiation. Also, we determined similar results in CD8+ CAR-T cells **(Figure 14B)**. These data show that PHA stimulation was the pioneer differentiation agent rather than 41BB or CD28 costimulatory domains on the CAR constructs.

**Figure 14:**
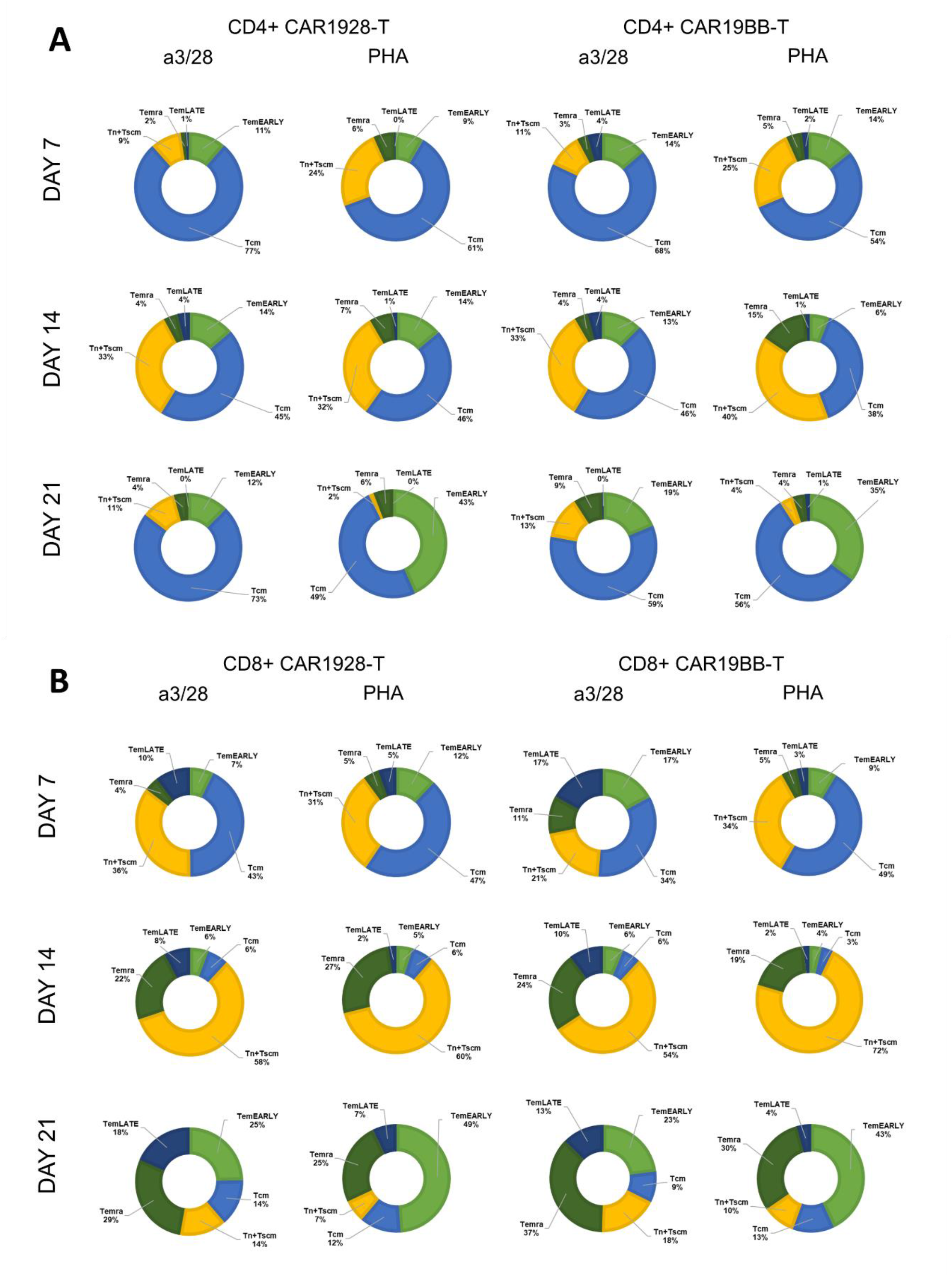
The pie-chart graphs showing T cell subpopulations expressing CAR(CD28) or CAR(4-1BB) in **A**. the CD4+ or **B**. the CD8+ T cell population after first and second activations on day 7, 14 and 21.

After characterization of T cell subpopulations upon activation process or CAR construct expression, we wanted to assess exhaustion biomarker expression on the cell populations. TIM3, PD1, and LAG3 expression were evaluated on day 14 and day 21 with the flow cytometry analysis **(Figure 15)**. No significant difference was observed in LAG3, TIM3, and PD1 MFI values of CD4+ CAR1928 or CAR19BB T cells at day 14 among anti-CD3/anti-CD28 and PHA-activated CD3+ T cells (**Figure 15B**). No significant difference was observed in LAG3, TIM3, and PD1 MFI values of CD8+ CAR1928 or CAR19BB T cells at day 14 among anti-CD3/anti-CD28 and PHA-activated CD3+ T cells (**Figure 15C**). A similar increase in LAG3, TIM3, and PD1 MFI values was observed in the comparison of total CD4+ T cells among anti-CD3/anti-CD28 and PHA-activated CD3+T cells. Therefore, no statistically significant difference was observed between PHA and anti-CD3/anti-CD28 (**Figure 15D**). When the LAG3, TIM3, and PD1 MFI values were compared in the comparison of total CD8+ T cells among anti-CD3/anti-CD28 and CD3+T cells activated with PHA, it was observed that LAG3 value decreased in PHA compared to anti-CD3/anti-CD28. However, there is no big difference **(Figure 15E)**. Second activation and post-culture in CD4+ or CD8+ CAR-T cells with anti-CD3/anti-CD28 or PHA No statistically significant difference was found in the expressions of exhaustion biomarkers LAG3, TIM3, and PD1 in CAR1928 and CAR19BB T cells compared to each other on day 7 (day 21 from the first activation) **(Figures 15F and 15G)**. When anti-CD3/anti-CD28 and PHA reactivations were evaluated in total CD4+ T cells, it was found that TIM3 and PD1 rates increased statistically significantly after PHA reactivity **(Figure 15H)**. When anti-CD3/anti-CD28 and PHA reactivations were evaluated in total CD8+ T cells, it was found that LAG3 and TIM3 rates increased statistically significantly after reagents with PHA (**Figure 15I**). These results show that exhaustion biomarkers increase more especially after reactivation with PHA. Here, we showed that the exhaustion marker levels were not significantly different in either PHA or aCD3/aCD28 activated T cells encoding CAR1928 or CAR19BB constructs.

**Figure 14:**
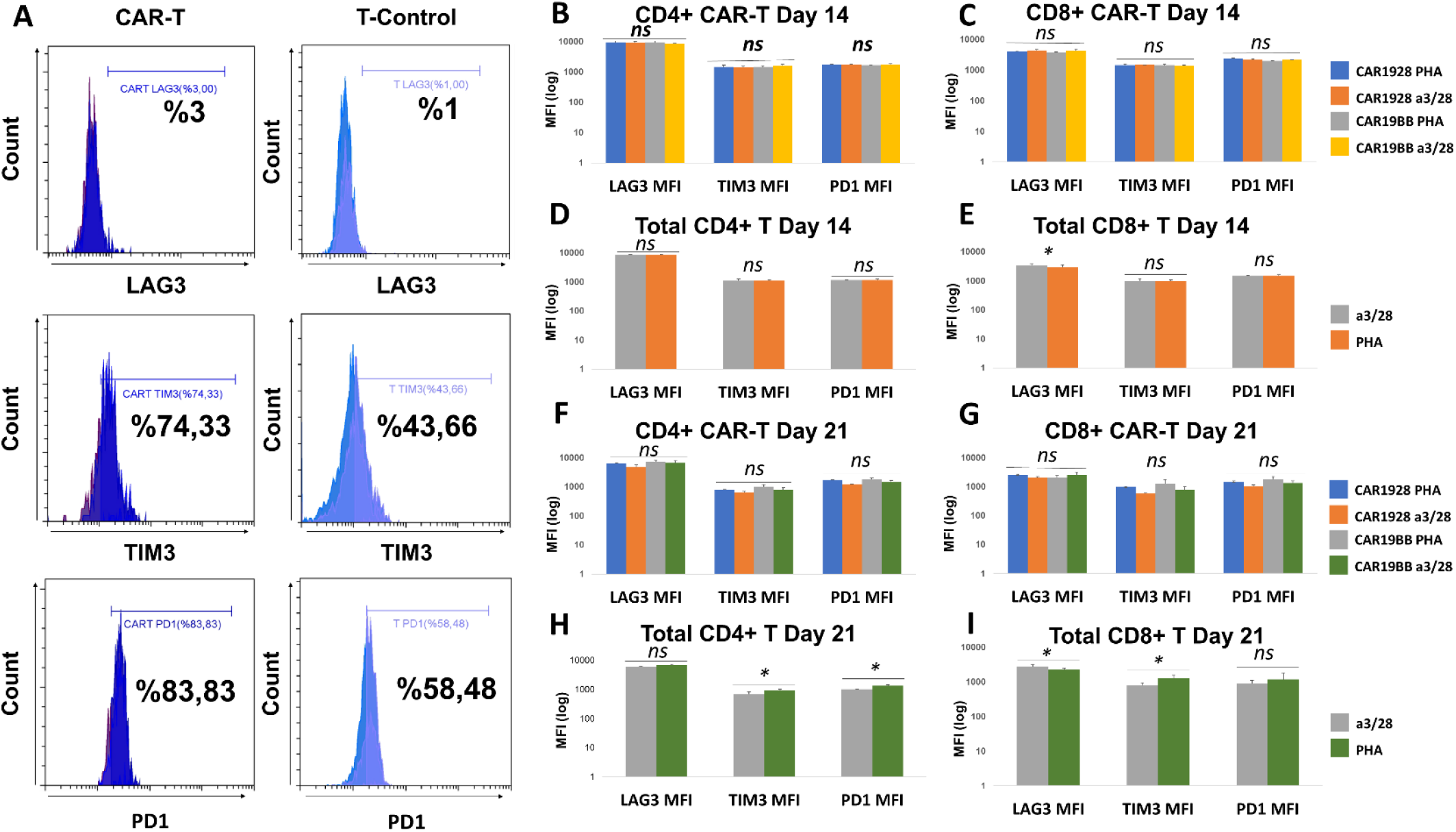
Exhaustion profile of PHA or aCD3/aCD28 activated CAR-T cells. **A**. Histograms showing LAG3, TIM3, and PD1 expressions of day 7 CAR-T cells activated with PHA after gating on CD3+, CD4+, CD8+. **B**. Comparison of MFI values of exhaustion biomarkers (LAG3, TIM3, PD1) of CD4+ CAR1928 or CAR19BB T cells at day 14 in CD3+ T cells activated with anti-CD3/anti-CD28 and PHA. **C**. Comparison of MFI values at Day 14 of exhaustion biomarkers (LAG3, TIM3, PD1) of CD8+ CAR1928 or CAR19BB T cells in CD3+ T cells activated with anti-CD3/anti-CD28 and PHA. **D**. Comparison of MFI values on day 14 of exhaustion biomarkers (LAG3, TIM3, PD1) of total CD4+T cells in CD3+T cells activated with anti-CD3/anti-CD28 and PHA. **E**. Comparison of MFI values on day 14 of exhaustion biomarkers (LAG3, TIM3, PD1) of total CD8+T cells in anti-CD3/anti-CD28 and PHA-activated CD3+ T cells. **F**. Comparison of MFI values on day 21 of exhaustion biomarkers (LAG3, TIM3, PD1) of total CD4+CAR-T cells in CD3+T cells activated with anti-CD3/anti-CD28 and PHA. **G**. Comparison of exhaustion biomarkers (LAG3, TIM3, PD1) 21st-day MFI values of total CD8+CAR-T cells in CD3+T cells activated with anti-CD3/anti-CD28 and PHA. **H**. Comparison of the MFI values of the exhaustion biomarkers (LAG3, TIM3, PD1) of total CD4+T cells (LAG3, TIM3, PD1) on Day 21 among CD3+T cells activated with anti-CD3/anti-CD28 and PHA. **I**. Comparison of exhaustion biomarkers of total CD8+T cells (LAG3, TIM3, PD1) on Day 21, among CD3+T cells activated with anti-CD3/anti-CD28 and PHA.

### Cytotoxic capacity of PHA or aCD3/aCD28 activated CAR-T cells

Next, we aimed to determine anti-cancer capacity of the CAR-T cells encoding CAR1928 or CAR19BB following activation and expansion with PHA or aCD3/aCD28 for 14 days. RAJI B cell lymphoma line were stained with aCD19 while CAR-T cells in CD3+ T cells were determined with aEGFR. To assess the activation of the cells, upregulation of CD25 and CD107a activation and degranulation markers were determined **(Figure 16)**. It has been observed that CAR19BB a3/28 has high killing proficiency on day 2, especially at 5:1 and 10:1. But by the 7th day, the kill rate has dropped to almost 0 **(Figure 16B)**. After the 7th day, it was observed that the rate of cytotoxicity increased to 100% again when given in combination with CAR1928. For CAR1928 a3/28, almost 100% anti-cancer effect was observed on the 2nd day, while it was observed that it killed 100% by the 7th day **(Figure 16B)**. A very low cytotoxicity rate was observed in CAR19BB PHA, even at a ratio of 10:1. However, even when CAR1928 CAR-T was produced by activating it with PHA, it was observed that it killed close to 100% on the 2nd day and 100% on the 7th day **(Figure 16B)**. Combining CAR1928 CAR-T, which is produced by activating with PHA, together with CAR19BB has also been observed to effectively kill RAJI. In the anti-cancer efficacy experiment with CAR-T cells produced by the second activation, RAJI cells became overpopulated on day 7 in all experimental groups **(Figure 16C)**. The anti-cancer activity was not successful in any CAR-T (PHA or α3/28 activated CAR19BB or CAR1928) cell co-culture experiment **(Figure 16C)**. This shows that CAR-T cells produced by the second activation have lost their anti-cancer activity to a large extent. Although it still seems to have anti-cancer activity when only CAR1928 is produced by activating T cells with α3/28, RAJI cells continue to proliferate on the 7th day **(Figure 16C)**. Therefore, it has been observed that PHA may not suppress the anti-tumor activity with CAR19BB and may show anti-tumor activity effectively with CAR1928.

**Figure 16:**
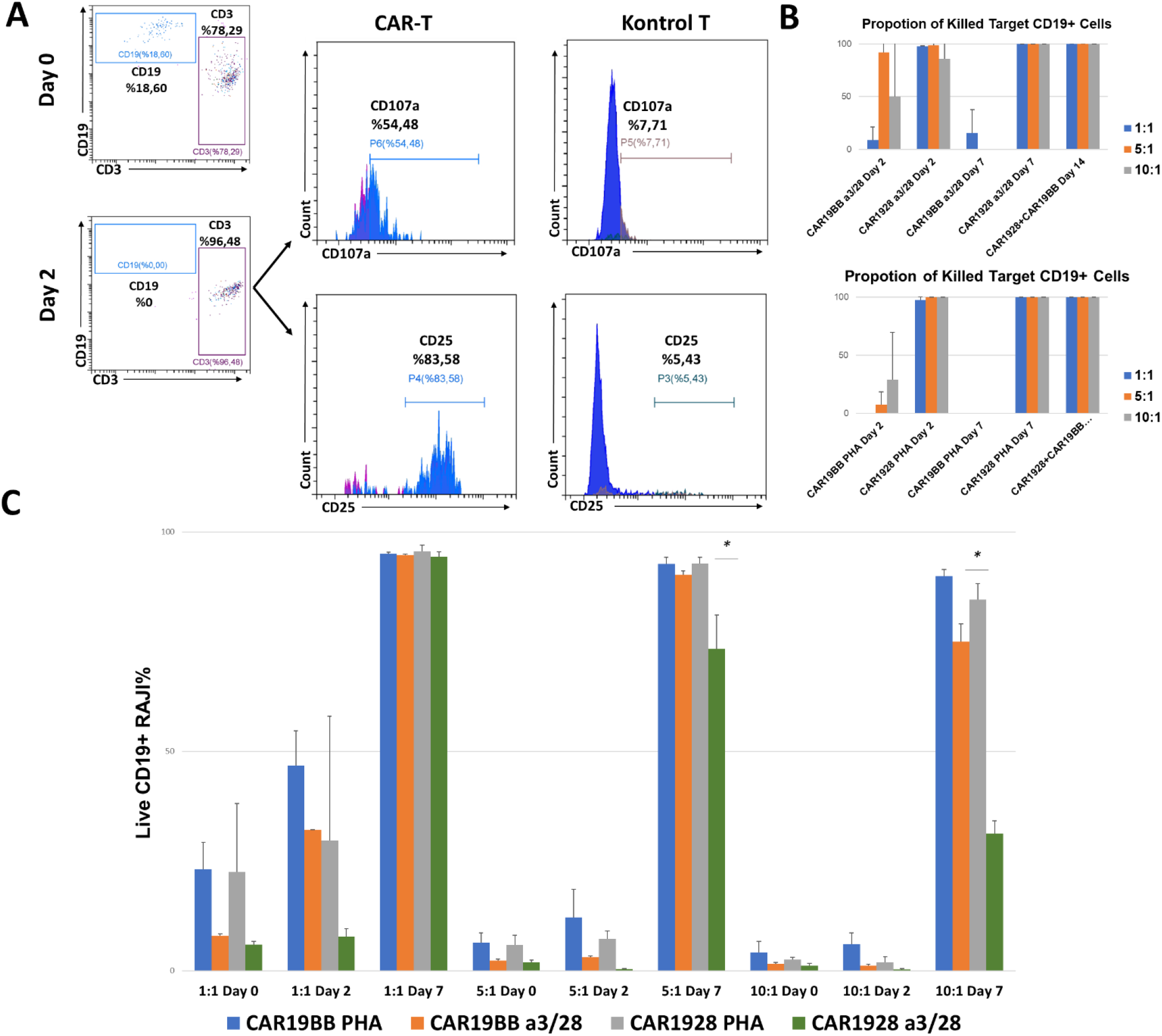
Cytotoxicity capacity of the CAR-T cells. **A**. Analysis of CD25 activation and CD107a cytotoxic de-granulation biomarkers of CAR-T cells and control T cells in CD3+ T cells by flow cytometry and death of CD19+ RAJI cells after 48 hours in CAR-T RAJI co-culture experiments. **B**. CD19+ RAJI Killing Assay Mortality Rates of CAR19BB and CAR1928 CAR-Ts produced by activating PHA and anti-CD3/anti-CD28. **C**. Bar graph showing the percentage of living CD19+ RAJI Cells, the anti-cancer activity with CAR-T cells produced by the second activation.

In the activation of CAR1928 T cells with both anti-CD3/anti-CD28 and PHA (1:1 CAR-T:RAJI), CD25 expression was observed to occur successfully for CD4+CAR-T and CD8+CAR-T. It was observed that CAR19BB exhibited a significant increase in activation by both anti-CD3/anti-CD28 and PHA compared to control CD4+T and control CD8+T. Compared to CD28, the rate of CD107a increased equally in CAR19BB **(Figures 17A and 17B)**. In the activation of CAR1928 with both anti-CD3/anti-CD28 and PHA (5:1 CAR-T:RAJI), CD25 expression was observed to occur successfully for CD4+CAR-T and CD8+CAR-T. Compared to CAR1928, it was observed that CAR19BB did not show effective CD25 activation by both anti-CD3/anti-CD28 and PHA. The rate of CD107a was also increased at a similar rate in CAR19BB compared to CD28. CD8+CAR-T CAR1928 PHA ratio was higher than the CD4+CAR-T CAR1928 PHA ratio **(Figures 17C and 17D)**. In the activation of CAR1928 with both anti-CD3/anti-CD28 and PHA (10:1 CAR-T:RAJI), CD25 expression was observed to occur successfully for CD4+CAR-T and CD8+CAR-T. Compared to CAR1928, it was observed that CAR19BB did not show effective CD25 activation by both anti-CD3/anti-CD28 and PHA. The rate of CD107a was higher in CD8+CAR-Ts compared to CD4+CAR-Ts. Although effective CD25 activation was not observed in CAR19BB CAR-T cells, similar activation was observed in CD107a cytotoxic degranulation with CAR1928 CAR-T cells **(Figures 17E and 17F)**.

**Figure17:**
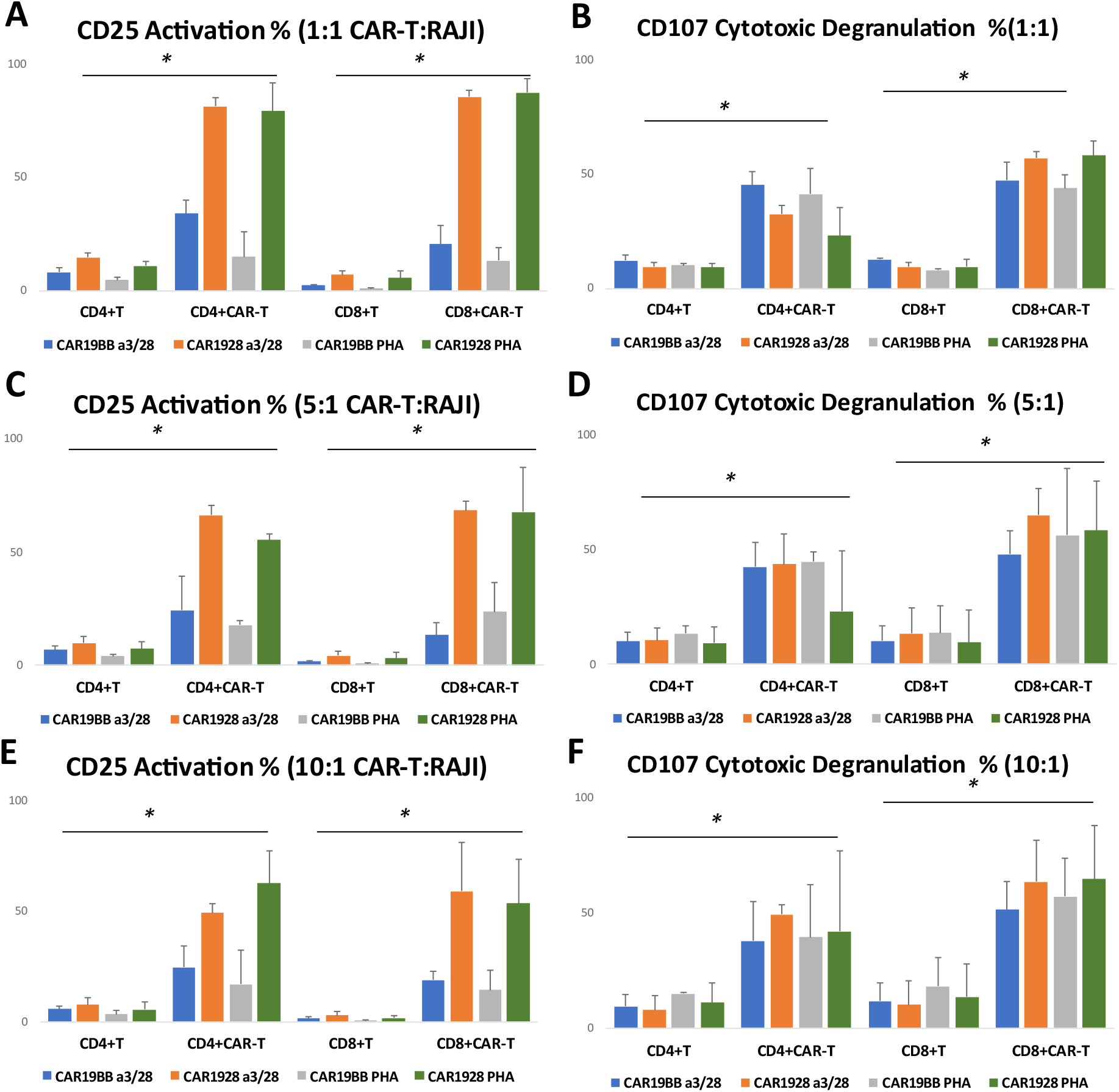
Activation capacity of CAR-T cells with CD25 and CD107a upregulation. Bar graph of **A**. CD25 activation and **B**. CD107a cytotoxic degranulation biomarkers of CAR-T cells and control T cells in CD3+T cells after 48 hours in 1:1 CAR-T: RAJI co-culture experiments. Bar graph of **C**. CD25 activation and **D**. CD107a cytotoxic de-granulation biomarkers of CAR-T cells and control T cells in CD3+T cells after 48 hours in 5:1 CAR-T: RAJI co-culture experiments. Bar graph of **E**. CD25 activation and **F**. CD107a cytotoxic de-granulation biomarkers of CAR-T cells and control T cells in CD3+T cells after 48 hours in 10:1 CAR-T: RAJI co-culture experiment.

## DISCUSSION

The time-dependent efficacy and stability of CAR-T cell applications, which is the most promising method in cancer treatments, needs to be improved. Currently available CAR-T cell applications use aCD3/aCD28 microbeads to activate cells. Here we report that PHA could be a good alternative for the activation and expansion of CAR-T cells to remain more stable and transform into memory cell groups after our results.

While the existing products in cellular immunotherapy are ineffective in creating complete remission, they are only short tools expressed separately in survival times. The main reason is that the existing immune cells carry low-activity receptors even if they are stimulated against cancer tissue. In this study, autologous T lymphocytes to be prepared by genetically altering CAR technology, which has been used in modern medicine, are provided to carry high-affinity receptors, and a product with more effective and stable results has been developed. For CAR-T cells to remain long-term and effective in *in vitro* and *in vivo* cancer models, it is aimed to provide these cells with a central memory feature (Tcm) and stem cell-like memory feature (Tscm). We have also developed a new alternative CAR-T manufacturing process to replace anti-CD3/anti-CD28 microbeads. And we have also shown that CAR expression can be increased at the desired rate by transduction after the reactivation. We aimed to increase the survival time and decrease the probability of cancer cell growth using PHA. When the proliferation ability of aCD3/aCD28 microbeads was examined, we observed that PHA significantly increased proliferation. Thus, it was observed that CAR-T cell production with PHA was similar to anti-CD3/anti-CD28 and had better production at some points. In this way, CAR-T cells will be able to multiply more quickly and easily before they are given to animals for clinical studies and this will provide a significant convenience in production. In both T cell types, the TemEARLY cell population was quite high in cells with PHA activation. This difference was also statistically significant. More TemEARLY cells mean there is a higher proportion of not exhausted yet memory cells in cells with PHA. TemEARLY cells possess a range of effector functions and are predominant in the target tissues. They are more likely to respond to tissue antigen reload than other T cell subpopulations. TemEARLY cells likely perpetuate autoimmune diseases due to their effector functions and relative longevity. Persistent antigen increases the pool of Tem cells, as demonstrated in studies of chronic infections; this would also be true in the context of autoimmune diseases where self-antigens persist (Devarajan et al., 2013).

The significant value of TemEARLY cells formed by activation with PHA shows that after CAR-T cell application, CAR-T cells remain in the patient’s blood for a longer time as memory cells and that it will be an important defense mechanism in case of relapsing cancer. Apart from the high number of TemEARLY cells, another important result of ours was that the Temra and TemLATE cell populations were less numerous in the cells with PHA. Memory cells are of great importance in terms of time-dependent efficacy and stability after CAR-T cell therapy. Because this makes it easier to analyze whether the cancer is overgrowth or cancer-free, depending on the number and effectiveness of memory cells. That’s why we have also tested, these CAR constructs compared to each other with different variations to analyze aCD3/aCD28 or PHA can make any significant differences depending on diverse CAR constructs. However, when we consider Total CD4+ and CD8+ T cell groups, we also found that memory cells can be preserved after the activation with PHA. Also showed us the cells are significantly farther from the exhausted profile. These results show that exhaustion biomarkers increase more especially after reactivation with PHA. It also explains that T cells expressing CAR1928 or CAR19BB in anti-cancer studies show higher anti-cancer capacity, especially with a3/28 reactivation. These results show that reactivation is not a preferred method.

Combining CAR1928 CAR-T, which is produced by activating with PHA, together with CAR19BB has also been observed to effectively kill the RAJI cell line. Therefore, it has been observed that PHA may not suppress the anti-tumor activity very much and may show anti-tumor activity effectively. This shows that CAR-T cells produced by the second activation have lost their anti-cancer activity to a large extent. It was investigated whether anti-CD3/anti-CD28 microbeads increase the Tcm and Tscm population or activation with PHA, a lectin that binds to the membranes of T cells and increases metabolic activity and cell division, would increase this rate. We have also developed a new alternative CAR-T manufacturing process to replace anti-CD3/anti-CD28 microbeads. This is the first result showing that T cells can be transduced by adding CAR lentiviruses in the first activation with PHA compared to anti-CD3/anti-CD28.

CAR-T cell production with PHA had no adverse effects on cell viability compared to aCD3/aCD28. Thus, the use of PHA as an alternative to aCD3/aCD28 in CAR-T cell production has also contributed to the literature. We have also shown that CAR expression can be increased at the desired rate by transduction after the reactivation. Based on our initial results that the combined use of CAR19BB and CAR1928 CAR-T cells instead of CAR-T cell production with the second reactivation may have permanent and long-term anti-cancer efficacy, combined CAR-T co-culture tests with CAR-T cells to be regenerated with independent donors is planned as well. Thus, we plan to take advantage of both the CAR19BB-mediated memory T cell ratio and use the high anti-cancer capacity of CAR1928-mediated T cells from the very first moment. In conclusion, we have shown that CAR-T cell production with PHA has a similar/better proliferation capacity than CAR-T cell proliferation obtained in anti-CD3/anti-CD28 activation. Following this, it was decided to compare the profiles of anti-cancer activity and Tcm-Tscm ratios of CAR-T cells activated with PHA to anti-CD3/anti-CD28 activated CAR-T cells.

